# Antibody background in ChIP-seq skews estimates of cohesin positioning by CTCF barriers

**DOI:** 10.64898/2026.02.26.708306

**Authors:** Yao Xiao, Erika C. Anderson, Hadi Rahmaninejad, Elphège P. Nora, Geoffrey Fudenberg

## Abstract

Loop extruding cohesin complexes are positioned by CTCF barriers to generate locus-specific 3D genome folding patterns. Quantifying cohesin accumulation at CTCF sites is crucial for deriving insight into loop extrusion and its consequences. Here we confronted expectations from loop extrusion simulations with experimental data by developing a pipeline, ChIP-FRiP, and analysis framework to reliably quantify cohesin positioning. We used ChIP-FRiP to uniformly re-process 140 cohesin ChIP-seq datasets from 13 publicly available studies. This analysis revealed that non-specific antibody binding can skew measurements of cohesin positioning. To mitigate this bias, we developed a biochemical model of background ChIP-seq signal and a strategy to estimate and correct this background using spike-in ChIP-seq data and relative protein abundance before and after depletion. Our results establish a framework for comparative analysis, demonstrating that accurate background correction is requisite for interpreting the roles of cohesin cofactors in cohesin positioning.

## Introduction

To orchestrate genome folding in mammalian interphase, the loop extruding cohesin complex is blocked by the protein CTCF(1, 2). This leads to accumulation of cohesin at CTCF binding sites evident in ChIP-seq data(3–5). Simulations of loop extrusion provide clear predictions for how cohesin accumulates at CTCF sites based on barrier strength and extrusion dynamics (6–8). Still, whether available data for cohesin positioning can be used to infer the roles of individual cohesin complex subunits and cofactors in extrusion dynamics remains unknown.

Since ChIP-seq provides a genome-wide measurement of protein positioning(9, 10), comparing cohesin ChIP-seq before and after perturbations to cofactors could potentially illuminate their roles in modulating extrusion dynamics. For example, depleting the cohesin unloader WAPL could alter cohesin positioning by modulating the number of cohesin extruders, their residence time, or their interactions with CTCF. However, such inferences require a quantitative measure of cohesin positioning before and after perturbation. The fraction of reads in peaks (FRiP) for a given protein is widely used as a quality-control measure for ChIP-seq, including in the ENCODE guidelines(11). However, FRiP can also be used to quantify the fraction of reads for one protein in the peaks of another protein. For determining how effectively CTCF blocks cohesin in different scenarios, the FRiP of cohesin reads at CTCF peaks(12) provides a natural quantification (**Fig. 1a**).

**Figure 1.**
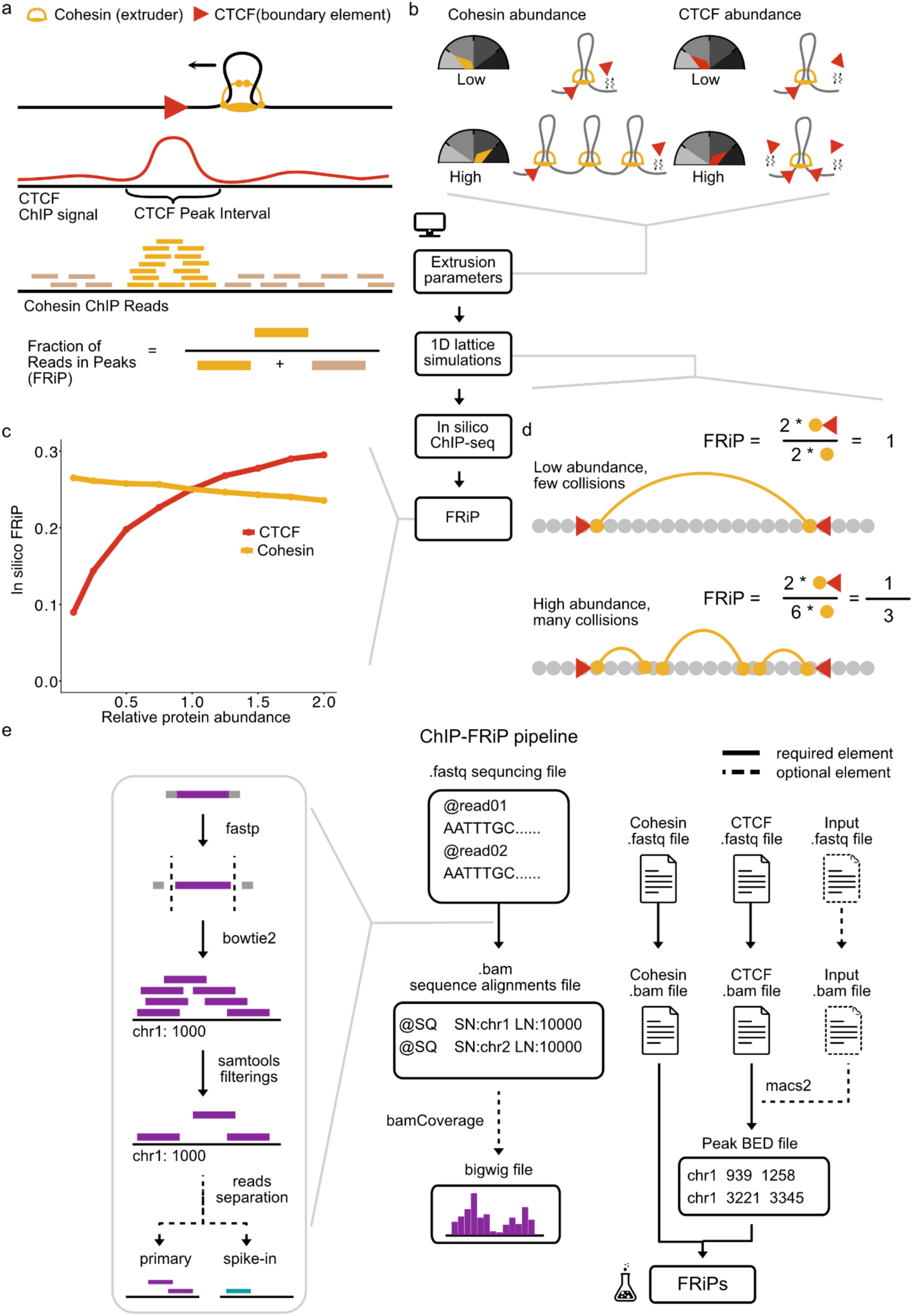
Cohesin positioning by CTCF quantified in simulations and experimental data. **a)** Cohesin extruders (black arrow shows movement) are blocked by CTCF barriers, leading to enrichment of cohesin ChIP-seq reads at CTCF peaks. The Fraction of Reads in Peaks (FRiP) quantifies the fraction of cohesin reads mapping within CTCF peak intervals. **b)** Simulations can be used to predict how modulating cohesin abundance or CTCF abundance impacts *in silico* ChIP-seq. **c)** Simulated FRiP gradually decreased as a function of cohesin abundance, and gradually increased as a function of CTCF abundance, from baseline abundance set to 1. **d)** Illustration of why increased extruder abundance decreases FRiP in the simulated system. The lattice used to represent position along the chromosome shown in grey, extruder leg positions shown as yellow dots with arcs indicating two legs of the same extruder complex, and red triangles represent CTCF barriers. **e)** The Snakemake-managed ChIP-FRiP pipeline can be used to quantify experimental cohesin positioning at CTCF sites. The pipeline accepts FASTQ files either from GEO or local datasets as input. Reads are then processed through quality control (fastp), alignment (Bowtie 2), and filtering (SAMtools) steps. Spike-in read separation and scaling factor computation is optionally applied when relevant. Peak calling is performed using MACS2. FRiPs are generated from the CTCF BED and cohesin BAM files using *bioframe*. Required pipeline elements are shown as solid lines, while optional components (spike-in read separation, ChIP input) are indicated by dashed lines.

Even though many publicly available ChIP-seq datasets profile cohesin and CTCF before and after perturbations to cohesin cofactors, computing FRiP of cohesin at CTCF in these datasets is not straightforward. First, computing FRiP requires aligned reads from cohesin ChIP-seq datasets, yet mapped reads are rarely deposited and hence must be regenerated from raw sequencing data. Second, FRiP requires peak calls from CTCF ChIP-seq datasets, which are more commonly deposited yet not uniformly processed. Third, available datasets differ in their choice of genome assembly, processing tools, parameters, antibody choice, sequencing depth, and ChIP-seq protocol, including whether input or spike-in controls are available. These challenges highlight two gaps for the field: 1. a pipeline capable of uniformly processing heterogeneous datasets to compute FRiP; 2. how technical covariates impact FRiP, and whether this alters biological interpretation.

Here we solve these challenges to enable uniform dataset processing and quantitative interpretation with biophysical models. First, we developed ChIP-FRiP, a Snakemake-based pipeline that reprocesses raw ChIP-seq FASTQ files into BAM and BED files that are required to calculate the fraction of cohesin reads within CTCF peak regions. Second, we conducted a meta-analysis of 13 publicly-available cohesin perturbation studies. We found substantial heterogeneity in FRiP across studies, even after perturbation of the same cohesin cofactors, as well as heterogeneity in untreated samples that depended on the tagged cofactor prior to any perturbation. Heterogeneity persisted despite considering biological and technical covariates such as cell types, target proteins and antibodies. Third, to understand how antibody binding and background noise can affect FRiP analysis, we coupled biophysical simulations of cohesin loop extrusion with a minimal biochemical model of ChIP-seq. This revealed that ChIP-seq background can invert the expected relationship between cohesin abundance and positioning. Our model shows that combining spike-in ChIP-seq and relative protein abundance before and after removal of the ChIP-ed protein can be used to estimate background noise and hence correct ChIP-seq quantification. Together, our results reveal previously neglected challenges in interpreting ChIP-seq data and highlight the importance of accurate background correction, which otherwise confounds biological interpretations - as we show for cohesin cofactors. More broadly, the framework we introduce streamlines quantitative analysis and sharpens the interpretation of ChIP-seq data across different biological and experimental contexts.

## Methods

### ChIP-FRiP Pipeline

This pipeline automates the processing of high-throughput sequencing data from ChIP-seq assays, and provides flexible usage for different situations: 1. w/o ChIP input 2. w/o spike-in.

For quality trimming and filtering, fastp (v0.23.4)(13) is used to process both single-end and paired-end reads, ensuring high-quality data for downstream analysis. Initial quality control assessments of raw and trimmed FASTQ files are conducted using FastQC (v0.11.5)(14).

Alignment of reads to reference genomes is performed with Bowtie 2 (v2.5.3), supporting both single-end and paired-end sequencing. The resulting SAM files are processed using SAMtools (v1.19) to filter low-quality alignments (-q), remove unmapped, secondary, or duplicate reads (-F 1804), and sort alignments by genomic coordinates with “samtools sort”. For paired-end data, mate-pair information is corrected using “samtools fixmate”, ensuring accurate downstream analyses. Duplicate reads are marked and removed using “samtools markdup”, reducing artifacts from library amplification.

Spike-in normalization is incorporated to ensure quantitative accuracy across samples generated from spike-in ChIP assay. The normalization procedure calculates the ratio of reads mapped to the primary genome versus the spike-in genome using “samtools idxstats”, and scaling factors are applied during bigWig file generation. BigWig files, both scaled and unscaled, are generated using bamCoverage from deepTools (v3.5.4.post1), enabling efficient visualization of sequencing data.

Peak calling is conducted with MACS2 (v2.2.9.1) to identify enriched regions in the genome, supporting both narrow and broad peaks, with or without input controls. Genome-specific analyses are further enabled by separating primary and spike-in genome reads using SAMtools and custom filtering steps, allowing for targeted processing and normalization.

The entire workflow is implemented using Snakemake (v7.32.4), ensuring reproducibility and scalability. This pipeline is designed to accommodate a range of experimental setups, including single-end and paired-end data, and can be easily configured with user-defined parameters such as quality thresholds, bin sizes, and scaling factors. By integrating spike-in normalization and robust quality control, this pipeline provides a user-friendly framework for processing ChIP-seq.

### Cohesin positioning meta-analysis

To quantify the relationship between cohesin positioning and CTCF binding, we employed the FRiP metric. To calculate FRiP, we aligned human and mouse sequence data to the hg38 and mm10 reference genomes, respectively. To ensure biological consistency, cohesin ChIP-seq reads were mapped specifically to CTCF peaks identified from the same biological replicate, considering only CTCF peaks identified in unperturbed samples. In datasets lacking explicit replicate pairing information, we utilized the median FRiP calculated across all available unperturbed CTCF peak sets. For studies with a single CTCF replicate, all cohesin FRiPs were computed relative to this single CTCF peak set. Notably, for the Arruda *et al.* and Kriz *et al.* datasets, which lacked matched CTCF ChIP-seq profiles, we utilized unperturbed CTCF peak sets from the Justice *et al.* dataset derived from the same cell type. Furthermore, to account for variations caused by covariates, we calculated the FRiP ratio between perturbations and unperturbed samples. These ratios were derived from samples matched by replicate and cohesin antibodies, using wild-type samples as the unperturbed reference when available. Code used for FRiP table generation and analyses were implemented in Python (v3.10) using the NumPy(15), Pandas(16), and Pysam(17) libraries.

### Quantifying covariates’ impacts on FRiP

To assess the explanatory power of each covariate, we compared their independent contributions to explaining FRiP (a proportion between 0 and 1) using beta regression. We fit separate beta regression models for each individual covariate and calculated BIC for each model using statsmodels.othermod.betareg.BetaModel (18) and its attribute (.bic). Categorical covariates were modeled with one parameter per category, and continuous covariates, such as total number of cohesin reads, were modeled with a single parameter. All models included an intercept term.

### Loop extrusion simulations for modeling cohesin number effects on FRiP

We adopted the dynamic barriers simulation framework for generating *in silico* FRiP (6). CTCF related parameters were set based on mESC occupancy data from Sönmezer et al.(19) and abundance from Cattoglio et al. (20). We computed abundance per megabase using an effective genome length of 3 * 2.7 Gb, assuming the average copy number during the cell cycle is 3. For CTCF, a total number of 217,200 and bound fraction around 49% yield an expected occupancy of 13 per Mb at baseline. For cohesin, following (21) we used 4 loaded cohesins per Mb (i.e. average separation between cohesins 250 kb) at baseline. We simulated loop extrusion on a 25 Mb sequence (25 replicas of 1Mb), both for computational efficiency and to maintain at least one cohesin extruder in the full simulation even at high depletion levels (note the number of cohesin extruders in simulations is set by the parameter “LEF_separation”, which controls the average distance between cohesins).

We sampled per-site CTCF occupancy parameters from Sönmezer et al. (19), and randomly positioned them on the 1 Mb sequence. CTCF occupancies were iteratively sampled until reaching the total target occupancy (13 per Mb). In the presented simulations, 38 CTCF site occupancies were sampled. To specify dynamic CTCF barriers, which require a bound time and unbound time, we assumed occupancy of 0.65 translates to a CTCF bound time of 9.87 minutes, as this provided good agreement for genome folding features (∼½ cohesin lifetime (6)). Using the corresponding unbound time at occupancy 0.65, we rescaled bound time and unbound time for the sampled CTCF occupancies. To compute simulated FRiP, we generated trajectories of 4000 steps. We discarded the first 25% of the trajectories to allow loop sizes to equilibrate. We then collected positions of cohesin legs, and computed the fraction of legs located at CTCF lattice sites using a window size of 1 (i.e., the CTCF lattice site plus the two flanking sites) to calculate FRiP. Simulation codes are publicly available at https://github.com/Yaoyx/ChIP-FRiP_manuscript_1d_simulation.

### Biochemical model of background noise in cohesin ChIP-seq

To investigate the effect of background reads, arising from non-specific antibody binding and/or other processes, we used an approach similar to (22) which we combined with ‘true’ reads for cohesin generated from the biophysical cohesin simulations described above.

We first defined the total ChIP reads as:

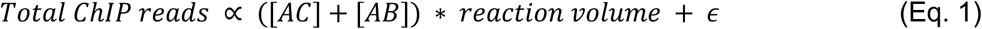

Where [*AC*] is the concentration of antibody bound to cohesin, [*AB*] is the concentration of antibody bound to background, and *ε* represents additional technical noise (e.g., wash efficiency for DNA fragments that are not bound with antibodies). We assumed that only chromatin-bound cohesin was available for antibody pulldown. With standard values for ChIP-seq experiments, we show below that [*AB*] is roughly independent of [*AC*] (or [*C*] or [*C^total^*]), and hence we could rewrite this equation as:

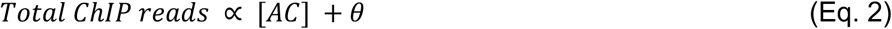

The full derivation is provided in Supplementary note, Equations S1-S8. Therefore, *θ* represents the total number of background reads. To adjust simulated FRiPs, we formulate this in terms of the fraction of background reads (*f*).

We investigated the relationship between *f* and cohesin abundance, assuming total background reads *θ* remains constant regardless of cohesin abundance. Hence, we calculated *θ* from a given background fraction *f_base_*, which represents the baseline background fraction. Defining *γ* as the relative cohesin abundance after a perturbation compared with its baseline abundance, the modified background fraction ( *f_mut_*) becomes:

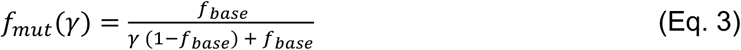

We note that [*AC*] cancels and the revised background fraction can be simply calculated from the initial background fraction. Finally, to study the impact of *f* on FRiP, we used revised background fractions to adjust FRiPs at different cohesin depletion or enrichment levels that were obtained through 1D loop extrusion simulation. In the simulation, each 1Mb replica is composed of 4000 lattice sites, each representing 250bp. We then control cohesin abundance by the average distance in monomers between two cohesins (LEF separation). The baseline LEF separation is set to 250kb. We then adjusted LEF separation to simulate different bound cohesin levels as ratios relative to the baseline LEF separation from 0.1 to 2. For all simulations, we use parameters: CTCF lifetime 780 sec, CTCF off-time 468 sec, LEF lifetime 1300 sec, LEF stalled lifetime 1300 sec, LEF birth rate 0.1, LEF pause rate 0.

### Estimating background fraction in ChIP-seq data

Due to the impact of background reads on FRiP, we therefore propose a way to estimate background fraction. This requires spike-in ChIP datasets that quantify the spike-in scaled number of reads for unperturbed (*R_UT_* ) and depleted (*Rdep* ) samples. We start by writing *Rdep* in terms of background reads and remaining cohesin reads:

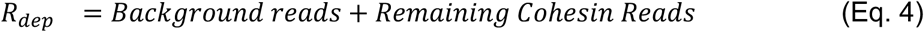

Using the condition-specific background fraction after depletion (Equation 3) and relative cohesin abundance (*γ*), we solve for the background fraction of the unperturbed sample (*f_UT_*) (see Equations S11-S12 for full derivation):

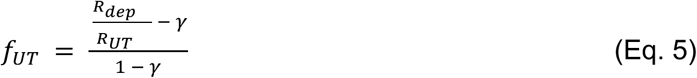

### Denoising FRiP using the estimated background fraction

Since FRiP is the ratio between reads in peaks and total reads, we need to remove background reads from both the numerator and the denominator to obtain denoised FRiP. We can define the denoised FRiP as follows:

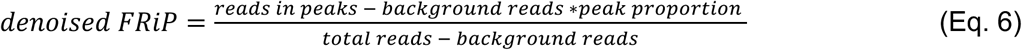

By substituting into Equation 6 and simplifying, we obtain:

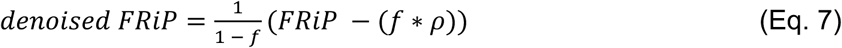

where *f* indicates the background fraction, and *ρ* indicates the proportion of the genome in peaks. We can compute *ρ* by dividing the number of basepairs in defined CTCF peak regions by the total number of basepairs in the genome. Either *f_UT_* estimated by Equation 5, or the condition-specific *f* obtained through Equation 3 can be substituted into Equation 7 to calculate the denoised FRiP.

## Results

### Simulations predict decreasing FRiP with increasing numbers of cohesins

Loop extrusion simulations provide clear expectations for how cohesin would accumulate at CTCF barriers depending on the kinetic parameters of cohesin extrusion (**Fig. 1b)**. Therefore, we first simulated how cohesin or CTCF depletion would impact the fraction of cohesin reads at CTCF peaks (hereafter FRiP). We used simulations of cohesin loop extrusion with dynamic CTCF barriers(6) and extrusion parameters chosen based on measurements in mouse embryonic stem cells (mESCs, **Methods**). These simulations model loop extrusion along a 1D lattice representing 25 Mb of chromatin at 250 bp resolution. Each cohesin is composed of two “legs” whose positions indicate the bases of the extruded loop (**Methods**). Positions of each leg are then aggregated across multiple simulations and trajectories to generate *in silico* ChIP-seq profiles, from which FRiP can be computed.

By running simulations either with variable numbers of cohesin extruders or with variable occupancy of CTCF barriers, we found two contrasting behaviors (**Fig. 1c**). First, as we decreased the average CTCF occupancy at barriers, the FRiP also decreased. This is intuitive, as an unbound CTCF site will not block cohesin, hence resulting in a loss of cohesin enrichment at CTCF sites. In contrast, as we decreased the number of cohesin extruders, FRiP at CTCF barriers gradually increased. Models attribute this behavior to reduced cohesin traffic: with fewer cohesins, each individual cohesin is less likely to collide with other cohesins, and hence more likely to arrive unimpeded at a CTCF barrier, increasing FRiP (**Fig. 1b**). Conversely, increased cohesin abundance leads to more collisions, resulting in fewer cohesins arriving at CTCF sites and lower FRiP.

### ChIP-FRiP pipeline from FASTA to FRiP

Having defined how FRiP is expected to vary upon changes in cohesin abundance or CTCF barrier activity, we next sought to compute cohesin FRiP in a uniform manner across available ChIP-seq datasets. To design a concise, reproducible, and user-friendly pipeline, we used the following design principles:

1. Employ a widely-used workflow manager.
2. Use widely-used tools for each data processing stage.
3. Curate default tool parameters from the ChIP-seq processing literature.
4. Support processing starting from fastq files, as these are available uniformly via SRA.
5. Support ChIP-seq data generated with multiple protocol variations.
6. Support arbitrary reference genomes and spike-in genomes.
7. Provide documentation and example configuration files on GitHub.

The major steps of our pipeline are: alignment, filtering, peak calling, and FRiP quantification (**Fig. 1d**), all provided as a Snakemake(23) script. For alignment, we use Bowtie 2(24). For filtering, we use SAMtools(25) to remove low mapq reads and duplicates. For peak calling we use MACS2(26). Finally, for FRiP computation we use bioframe(27). In addition to the main pipeline, we also provide utility Python scripts with a command-line interface (CLI) for automated metadata extraction from Gene Expression Omnibus (GEO) datasets using ffq(28), and systematic FRiP table generation to facilitate downstream comparative analyses. Code and documentation are publicly available at: https://github.com/Fudenberg-Research-Group/ChIP-FRiP.

ChIP-FRiP supports datasets generated with or without input controls and with or without spike-in normalization. Unlike existing pipelines optimized primarily for general ChIP-seq processing or specific protocol configurations(11, 29, 30), ChIP-FRiP was designed to support systematic FRiP quantification across diverse public datasets (**Table S1**). We validated the pipeline by comparing its output to Spikeflow on a spike-in control sample and to chipseq on a non-spike-in sample, obtaining nearly perfect correlation in both cases (**Fig. S1**). Although motivated by cohesin positioning at CTCF barriers, the pipeline is broadly applicable to any analysis requiring quantification of reads over defined genomic intervals.

### Meta-analysis of how cohesin perturbations impact experimental FRiP

We applied ChIP-FRiP to 13 studies with 140 publicly available ChIP-seq datasets for 16 distinct perturbations to key cohesin cofactors in mouse and human cell lines(31–43) (**Fig. S2, Table S2**). FRiPs for cohesin datasets before and after perturbations were computed relative to CTCF peaks from unperturbed cells. We use the term unperturbed to refer to both ‘wild-type’ and control datasets prior to explicit perturbations, where control datasets include cell lines where cohesin or cofactors have been tagged with degrons.

Despite uniform processing (**Fig. 2a**), FRiP for unperturbed datasets displayed substantial variation, ranging from 5% to 25% of cohesin reads in CTCF peaks across studies (**Fig. 2b**). We thus wondered which biological and technical covariates influenced FRiP, including: cell type, antibody type, the total number of cohesin reads, and whether a spike-in was included during library preparation (**Fig. 2c-f**, also **Fig. S3**). We determined the relative importance of each covariate by fitting a Beta regression model and quantifying the respective Bayesian Information Criterion (BIC) score (**Methods, Table 1**). Cohesin antibody type was least informative by BIC, reflecting the high FRiP variation even for identical antibodies. The total number of basepairs in CTCF peaks and the cell type were the most informative by BIC. While intriguing, it is challenging to ascertain if differences between cell types are driven by biology, due to the differences in protocols and other covariates.

**Figure 2.**
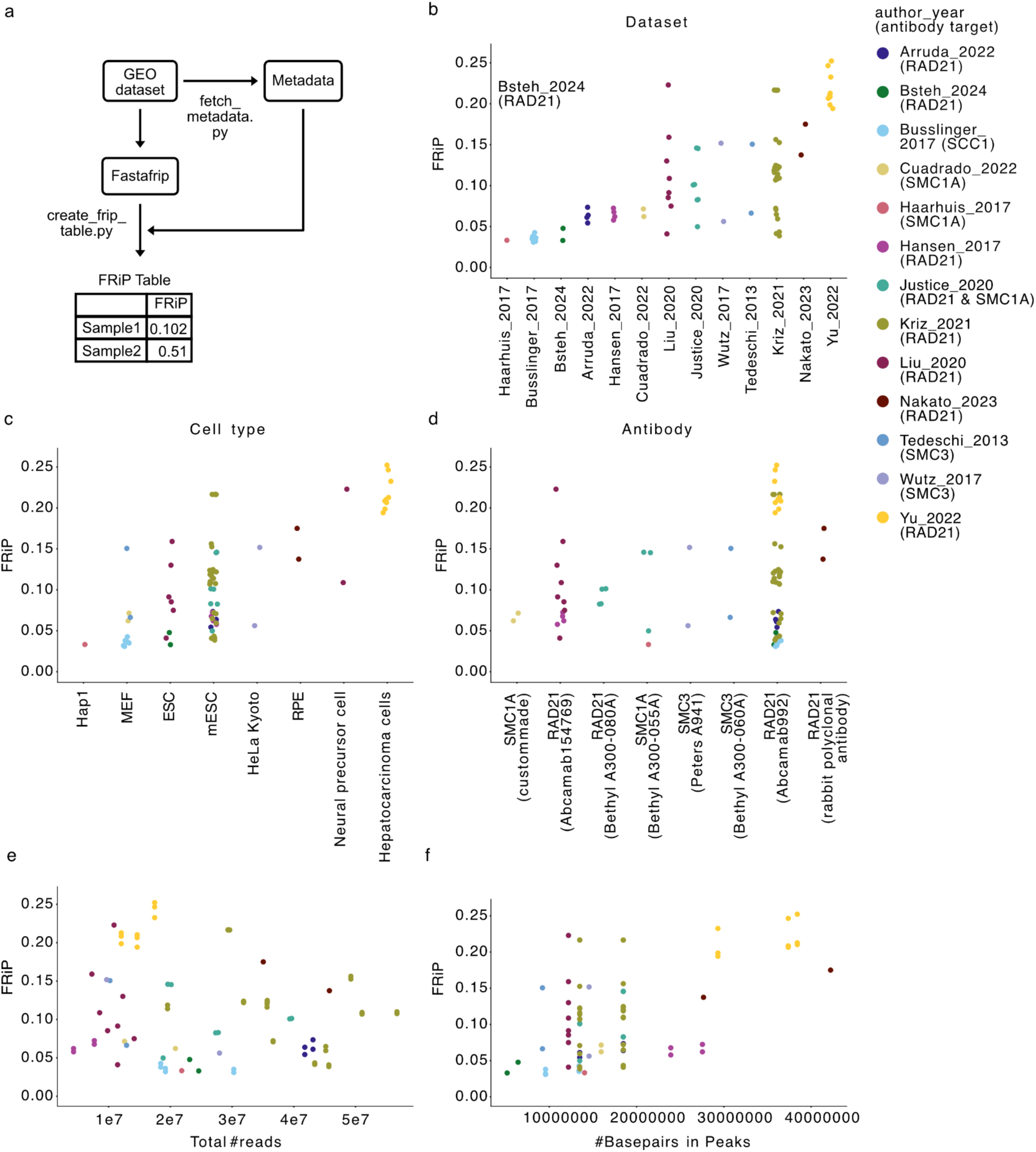
Cohesin positioning prior to perturbation shows substantial variation across biological and technical covariates. **a)** outline of using ChIP-FRiP to create FRiP tables for meta-analysis with script names next to the relevant processing steps. **b)** FRiP in unperturbed samples, highlighting wide variation at baseline across studies. Points are colored by study (author-year). **c)** FRiPs by cell type. **d)** FRiPs by antibody target. Categories in panels b-d are sorted by median FRiP. **e)** FRiPs versus total read count. **f)** FRiP versus the total number of base pairs in CTCF peaks. No single covariate accounts for FRiP variation across unperturbed samples, and available experimental data precludes jointly controlling for covariates.

**Table 1.** Comparison of covariates using Bayesian Information Criterion (BIC). Table of Beta regression BIC for covariates fit one at a time, covariates are ranked by their BIC values. Lower BIC values indicate a better model fit relative to complexity.

| Covariate | BIC |
| --- | --- |
| Total number of basepairs in CTCF peaks | -235 |
| Cell type | -232 |
| Spike-in usage | -220 |
| The number of CTCF peaks | -215 |
| Total number of cohesin reads | -207 |
| CTCF antibody | -203 |
| Cohesin antibody | -182 |

We also investigated the impact of protein tagging on FRiP in otherwise unperturbed cells, as degron tags can reduce protein expression(44). For multiple cofactors, we found that tagging alone made FRiPs deviate from those of untagged cells prior to any perturbation (**Fig. S3c-e**). We therefore recommend that experiments performing ChIP after protein depletion also include an untagged control to account for degron leakiness.

Since the range of FRiP observed in unperturbed samples approached that of perturbation samples due to covariates mentioned above (**Fig. S4**), we considered the FRiP ratio between matched samples before and after perturbations to determine the impact of specific factors on cohesin positioning. Because the studies we analyzed used different depletion strategies (e.g., RNAi, inducible degrons, genetic knockout), we refer to depleted samples by prefixing the protein name with ‘d’. We observed a range of FRiP ratios across the perturbation conditions (from 0.1916 to 2.070, **Fig. 3, Fig. S5**).

**Figure 3.**
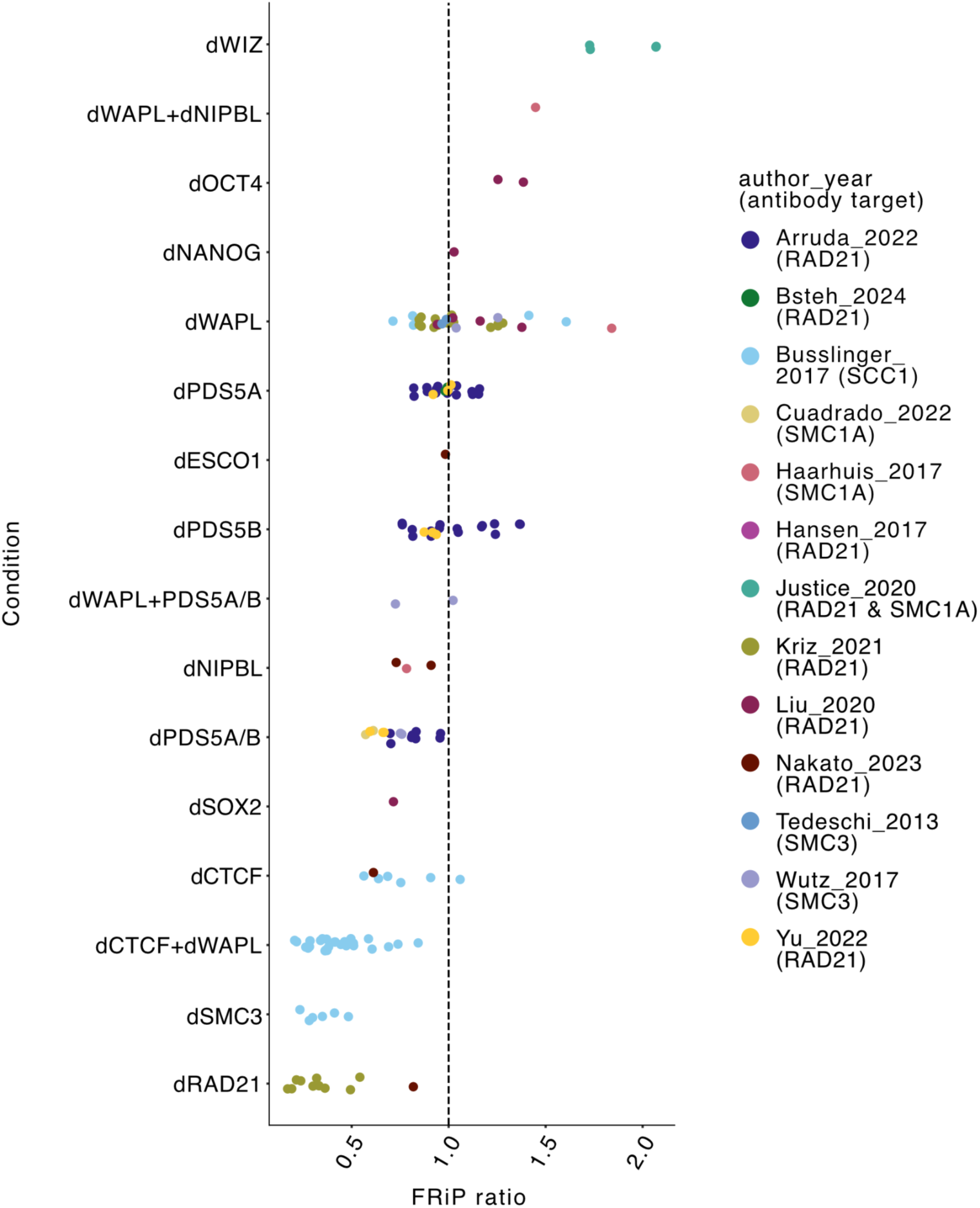
Cofactor perturbations alter apparent cohesin positioning. Ratio of perturbed to unperturbed FRiP for indicated cofactors and studies. FRiP ratios were computed for samples with matching replicate information. If there were multiple unperturbed samples, or replicate information was not available, we used the median unperturbed FRiP to compute the ratio. Each dot represents one computed ratio. Ratios above 1.0 indicate increased cohesin accumulation at CTCF peaks after perturbation. The “d” prefix denotes depletion of the corresponding cofactor.

For many perturbations, while we lacked specific predictions from simulations, we observed FRiP ratios that deviated from one. Depletion of WIZ, whose ChIP-seq peaks overlap substantially with those of CTCF(31), had the most substantial increase in FRiP ratio. This observation was surprising given that Hi-C following WIZ depletion did not display as many salient changes as after other cofactor perturbations, which we found impact FRiP less. For example, while WAPL depletion (dWAPL) greatly increases Hi-C contact frequencies between distant CTCF sites, it only mildly increased FRiP on average, with substantial variation between studies(32). Surprisingly, the combination of dCTCF and dWAPL decreased the FRiP further than dCTCF alone(38). Individual depletion of PDS5A or PDS5B resulted in a modest decrease in the FRiP ratio, while simultaneous depletion of both PDS5A and PDS5B produced a more dramatic reduction in the FRiP ratio, which was consistent across all four studies with dual depletion of PDS5A and PDS5B(33, 35, 39, 43). This aligns with hypothesized synergistic roles of these paralogous proteins in cohesin positioning(43). Other perturbations included depletion of ESCO1(36) and depletion of transcription factors OCT4, NANOG, and SOX2(42), though lack of replicates makes their impact on cohesin positioning uncertain.

For cohesin and CTCF, we had strong expectations from our simulations for how FRiP would be altered by their depletion (**Fig. 1b,c**). Consistent with our simulations, CTCF depletion (dCTCF) samples exhibited FRiP ratios below 1 (**Fig. 1b,c**). This agrees with the known role for CTCF barriers in positioning cohesin extruders, and previous observations of reduced FRiP after CTCF depletion(12). In contrast, we were surprised to observe very low FRiP ratios for SMC3 depletion (dSMC3) and RAD21 depletion (dRAD21), as decreased FRiP ratios after lower cohesin abundance diverged from the expectation from simulations (**Fig. 1**), prompting us to search for factors that could explain this behavior.

### Background reads in ChIP-seq can invert FRiP trends after cohesin perturbations

While simulations predicted that FRiP would increase with fewer cohesins, in experiments both dRAD21 and dSMC3 decreased the FRiP ratio. We hypothesized that this discrepancy may reflect non-negligible contributions of technical background to the ChIP-seq signal. Indeed, ChIP-seq background can emerge via non-specific antibody binding during immunoprecipitation (22). We found no evidence that background was driven primarily by problematic regions, as excluding regions in the ENCODE blacklist had negligible impact on FRiP (**Fig. S6**). Hence, if background reads could emerge from anywhere in the genome and given that CTCF peaks only represent a small part of the genome (∼0.8%), even using antibodies with low non-specific background affinities could substantially alter cohesin FRiP (**Fig. 4a**).

**Figure 4.**
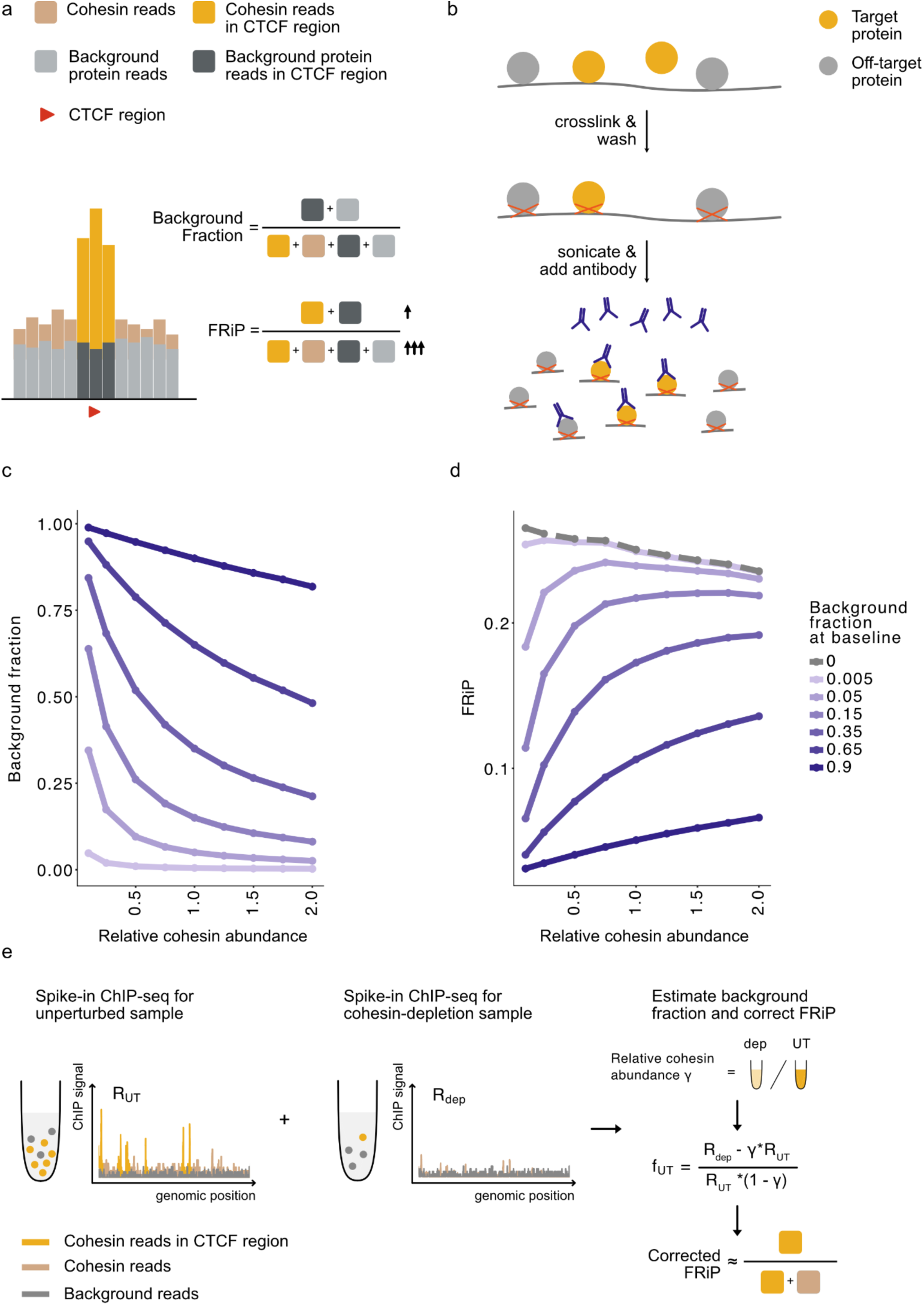
Biochemical model shows ChIP-seq background can invert the relationship between FRiP and cohesin abundance. **a)** Illustration of how uniformly distributed background reads decrease FRiP. **b)** Schematic of the uniform background model for ChIP-seq. The model assumes that antibodies capture target proteins directly bound to chromatin, with occasional binding to chromatin-bound background proteins. **c)** Background fraction does not scale linearly with cohesin abundance, and antibodies of varying specificity (modeled by different background fraction at baseline) elicit different scalings. **d)** corresponding simulated FRiP does not scale monotonously with cohesin abundance. Varying antibody specificity can invert whether FRiP increases or decreases with cohesin abundance. Dashed grey lines show simulations without background reads (as in Fig. 1c)**. e)** Illustration of how spike-in ChIP-seq combined with an orthogonal estimate of relative cohesin abundance before and after cohesin degradation can be used to estimate background fraction and correct FRiP. R = scaled number of ChIP-seq reads; UT = unperturbed; dep = depleted; γ = relative cohesin abundance.

To explore the effect of ChIP background, we developed a minimalistic biochemical model (**Fig. 4b)** that determines how background reads scale with cohesin abundance, and how this in turn alters FRiP (**Methods**). Our uniform background model assumes: 1. equilibrium binding kinetics for antibody, cohesin, and background proteins; 2. sufficient antibody abundance to capture all available cohesin; 3. the number of background-bound antibodies is much lower than the total number of antibodies; 4. Roughly uniform distribution of background targets across the genome.

Using this biochemical model, we computed FRiP as a function of cohesin abundance across a range of background levels. As expected, as cohesin abundance decreases, the fraction of background reads increases (**Fig. 4c**). Yet, this trend was not linear, and this nonlinearity was most pronounced for antibodies with the lowest background. We then investigated how these non-trivial background contributions modify the relationship between FRiP and cohesin abundance (**Fig. 4d**). In simulations without background, FRiP decreased with increasing cohesin abundance; however, introducing sufficient background reversed this relationship. When the background fraction exceeded ∼0.35, FRiP increased monotonically with cohesin abundance. There, increased signal from additional cohesin epitopes outweighed the negative effect of collisions on FRiP. These simulations further emphasize that FRiP varies quantitatively with cohesin abundance.

We considered whether other sources of background, such as proximal crosslinking of unbound cohesin (45–47), could also account for how FRiP decreases in experiments lowering cohesin abundance. If we assumed that all background reads originated from such proximal crosslinking, we observed that the monotonous scaling between FRiP and cohesin abundance was preserved (**Fig. S8b**). This occurred because the background for the proximal crosslinking model scaled with cohesin abundance, in contrast with the non-specific antibody binding model (**Fig. 4d**). Thus, a background source that is independent of cohesin abundance, such as non-specific antibody binding, is required to explain the reduction in FRiP after cohesin depletion observed experimentally.

Together, our minimal biochemical model provides a concise explanation for why FRiP can decrease after cohesin depletion in experimental conditions where antibodies are not entirely specific.

### Correcting for ChIP-seq background reads

Given the confounding effect of background reads on quantitative estimates in ChIP-seq, including FRiP, we considered how their impact could be mitigated in future experiments. Analysis of the uniform background model (**Methods**) suggests a practical strategy to correct FRiP for background by combining: (i) spike-in ChIP-seq before and after degradation of the target protein, and (ii) an orthogonal quantitative estimate of the relative abundance of the target protein after degradation (**Fig. 4e, S7a; Equation 5**). When complete removal by genetic knockout is not feasible, near-complete depletion through degrons or RNAi must be coupled to precise estimation of the leftover quantity, e.g. using a Halotag for quantitation(48). Indeed, our analyses show that failing to account for even low leftover levels markedly inflates background estimation and undermines the correction (**Fig. S7b**). In particular, under partial depletion but high antibody specificity, relying solely on the ratio of spike-in calibrated reads before and after depletion substantially overestimates background. However, as complete depletion is approached, we find that a simple ratio of spike-in calibrated reads performs well for estimating background, as recently applied for cohesin RAD21(49).

## Discussion

In this study, we developed and employed the ChIP-FRiP pipeline to determine how cohesin is positioned by CTCF across multiple perturbation studies. By consistently quantifying FRiP for cohesin reads in CTCF peaks, our meta-analysis revealed potential challenges to cross-study comparisons, both from biological and technical covariates. Indeed, cofactor tagging alone led to reproducible differences in apparent positioning. Perturbations of the same cohesin cofactor often displayed wide ranges of values, particularly for WAPL depletion. In loop extrusion simulations, we found that FRiP is expected to decrease as cohesin abundance increases, due to increased cohesin-cohesin collisions. However, our biochemical modeling revealed that antibody-related background noise can invert this relationship, highlighting how technical biases can impact biological interpretations of cohesin positioning. Analysis of our biochemical model suggests a strategy to estimate and correct for the impact of uniformly distributed background reads on FRiP using spike-in ChIP-seq and a quantitative measurement of target protein abundance. Together our results highlight several experimental challenges that must be addressed to understand precise roles of individual regulators of cohesin positioning.

Our combination of cohesin loop extrusion simulations and biochemical modeling provide intuition for how cohesin positioning can be influenced by experimental biases and covariates. First, our model of antibody-related background noise can explain why FRiP decreases after reducing the abundance of the core cohesin subunits RAD21 or SMC3. Second, the simulation results caution against simple mechanistic interpretations of experimental perturbations. For instance, how much WAPL depletion increases cohesin abundance at CTCF peaks emerges from the combination of multiple opposing effects: (i) increased cohesin lifetime allows more cohesins to translocate to CTCF peaks, increasing FRiP; (ii) greater number of loaded cohesins causes more collisions, decreasing FRiP; (iii) increased abundance of loaded cohesins reduces the background fraction, increasing FRiP. Similarly, reduced fraction of cohesins at CTCF peaks following NIPBL depletion might arise either due to lower extrusion processivity or, technically, from decreased abundance of cohesin epitopes on chromatin – not simply due to fewer cohesins being loaded on chromatin. Finally, it will be interesting to confirm in other cell types and determine which mechanisms contribute to the prominent impact of WIZ on cohesin positioning highlighted by our metanalysis.

More broadly, our results call for cautious interpretation of differences in cohesin positioning across studies, as they may reflect unmatched covariates (e.g. differences in absolute cohesin levels between cell types or use of antibodies of varying specificity), rather than mechanistic differences in the function of the perturbed cofactors. Importantly, our study highlights ways to mitigate these limitations (e.g., profiling tagged unperturbed cells, precisely evaluating antibody specificity, using spike-in calibration, and quantitatively measuring bound cohesin levels) and provides computational methods to support mechanistic study of cohesin dynamics in cells.

We acknowledge limitations of our current approach that warrant future investigation. First, our meta-analysis assumed that CTCF peak positions were largely unaltered by cohesin perturbations, since CTCF binding was not systematically profiled after each cohesin perturbation. Second, available studies have limited overlap in the choice of cell types, antibodies, protocols, and perturbation strategies making it challenging to control for covariates when assessing the impact of cohesin regulators. Third, our minimalistic loop extrusion simulations model cohesin positioning based only on CTCF while neglecting potential contributions from cohesin cofactors. Fourth, our biochemical model relies on four key assumptions: equilibrium kinetics for proteins, sufficient antibody abundance to capture all cohesin, background signal is weaker than the cohesin signal, and background has a roughly uniform distribution across the genome. These assumptions may not fully reflect cohesin ChIP assay conditions used in practice. Finally, our background fraction estimation relies on the assumption that the total abundance of accessible epitopes (generating both target and background reads) is similar across samples prior to depletion. This presumes that both the mass of chromatin input and the accessibility of the target epitope are constant across experiments. In practice, however, due to technical differences in nuclear extraction efficiency, fixation, and purification, the effective recovery of protein-bound chromatin may vary between experiments, limiting the accuracy of this estimate.

## Code and Data Availability

The ChIP-FRiP pipeline is available at https://github.com/Fudenberg-Research-Group/ChIP-FRiP. All ChIP-seq datasets analyzed in this study are publicly available and GEO accession numbers are provided in Supplementary Table 2. Simulation codes are available at https://github.com/Yaoyx/ChIP-FRiP_manuscript_1d_simulation.

## Acknowledgements

The authors thank Lin Chen and Mark Chaisson for manuscript feedback.

