## Supplementary Materials for "Antibody background in ChIP-seq skews estimates of cohesin positioning by CTCF barriers"

### Supplemental Material for: Antibody background in ChIP-seq skews estimates of cohesin positioning by CTCF barriers

a

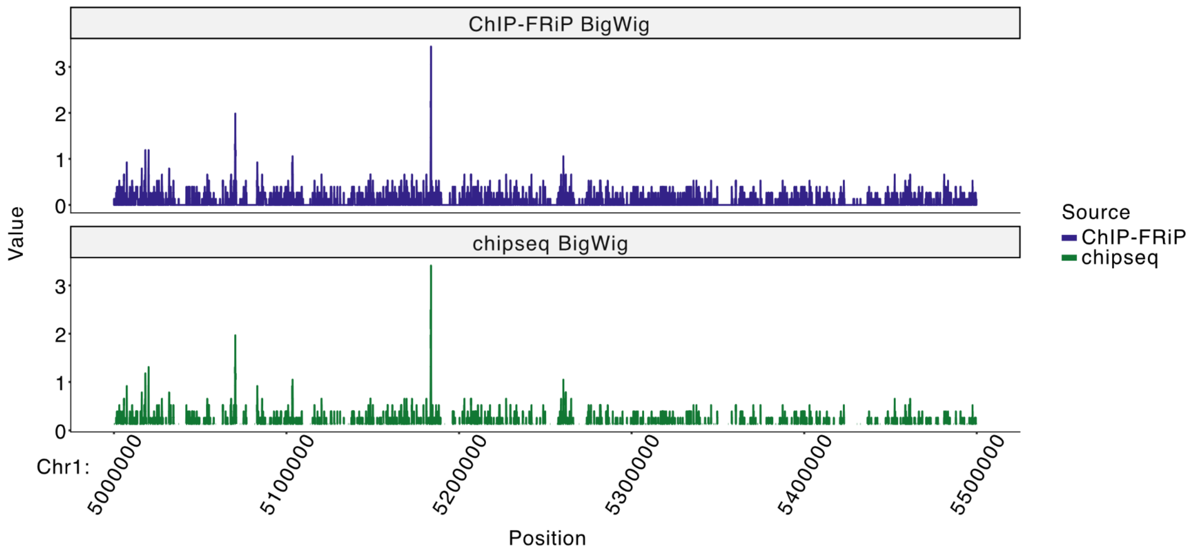


b

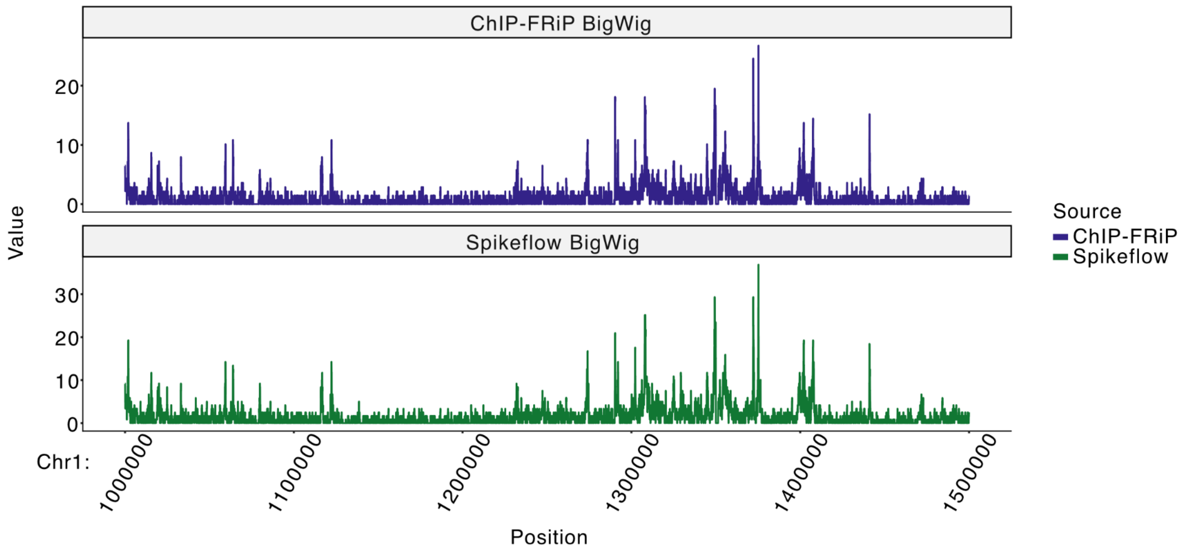


**Figure S1. ChIP-FRiP yields comparable results to other pipelines for both non-spike-in and spike-in ChIP-seq data. a)** Non-spike-in ChIP-seq data from mESCs (SRR5085155**(37)**): ChIP-FRiP is highly concordant with the CPM-normalized bigWig tracks output of the nf-core/chipseq pipeline**(30)** (Pearson's r = 0.98). **b)** Spike-in ChIP-seq in hepatocarcinoma cells(SRR20664875**(33)**): ChIP-FRiP is highly concordant with the raw counts output of the Spikeflow pipeline**(29)** (Pearson's r = 0.96). ChIP-FRiP has the additional option of directly applying the spike-in scaling factor to generate a normalized signal track.

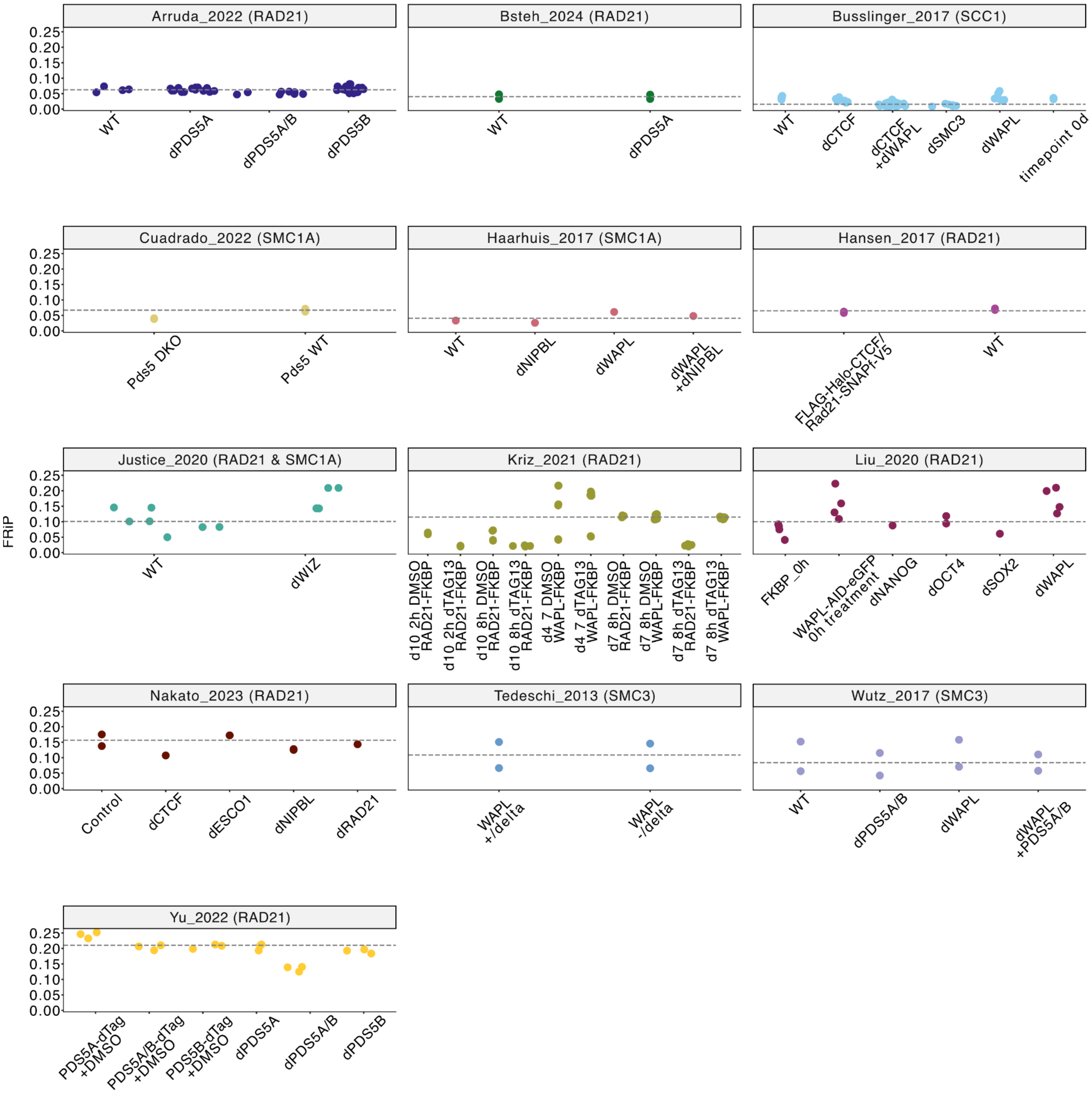


**Figure S2. FRiPs collected from 13 studies reveal a range of behaviors after cofactor perturbation**. Each point represents an individual cohesin ChIP-seq sample. Grey dashed lines indicate the median FRiP for unperturbed samples as provided in the metadata of each study.

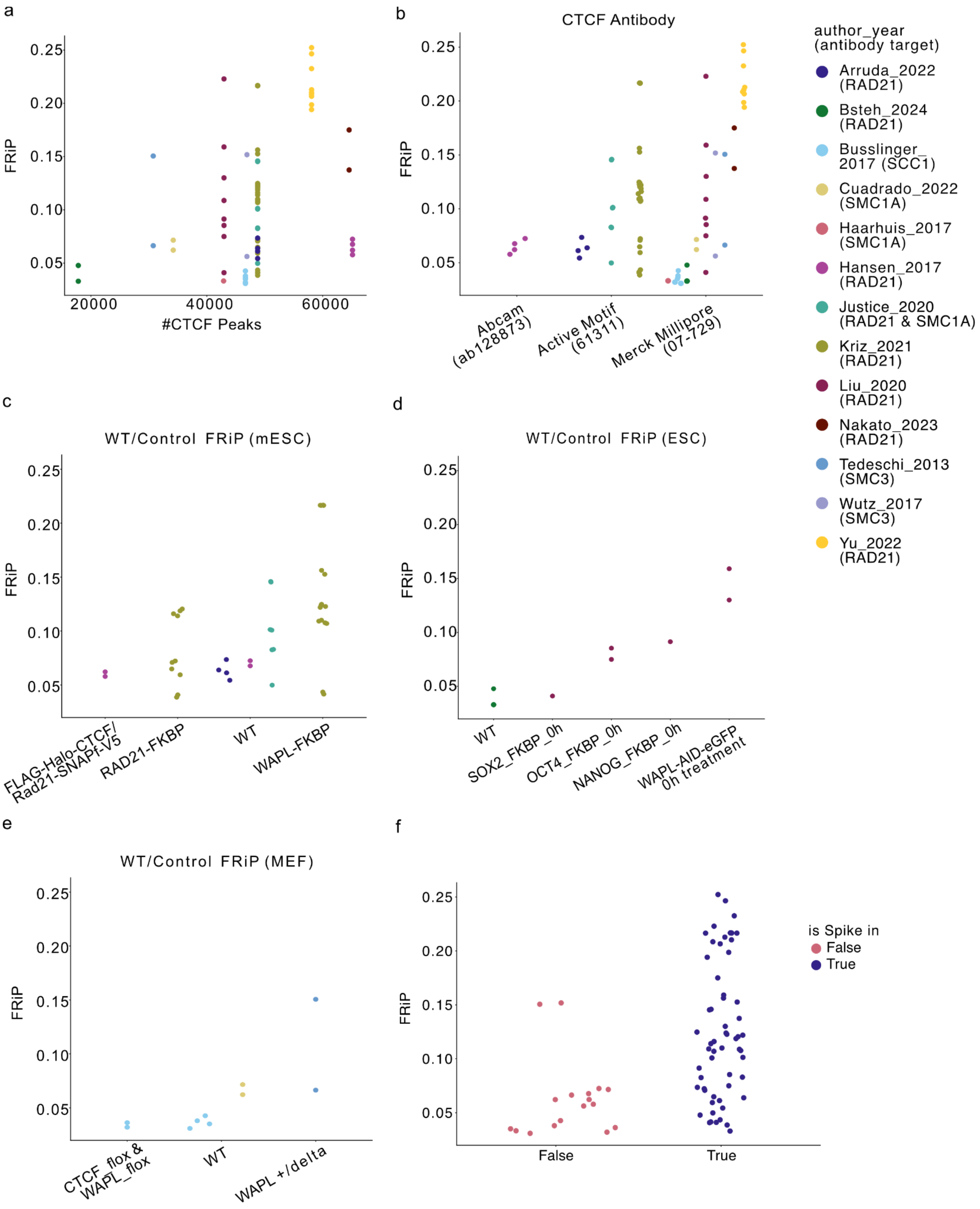


**Figure S3. Additional covariates that might affect FRiPs in unperturbed datasets. a)** FRiPs versus the number of identified CTCF peaks. **b)** FRiP versus CTCF antibody. **c)** FRiPs versus TAG type in mESC. **d)** FRiPs versus TAG type in ESC. **e)** FRiP versus TAG type in MEF cell. **f)** FRiP versus spike-in usage.

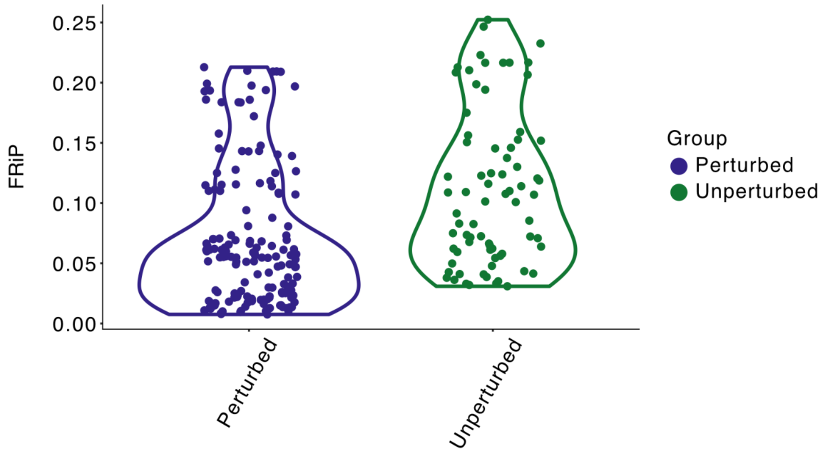


**Figure S4. Similar FRiP variance between unperturbed and perturbed conditions.** Each point represents FRiP computed for either a perturbed or unperturbed sample, aggregated across all studies. The wide dispersion of FRiP across unperturbed samples argues against simple comparisons of FRiPs across studies and conditions without normalization.

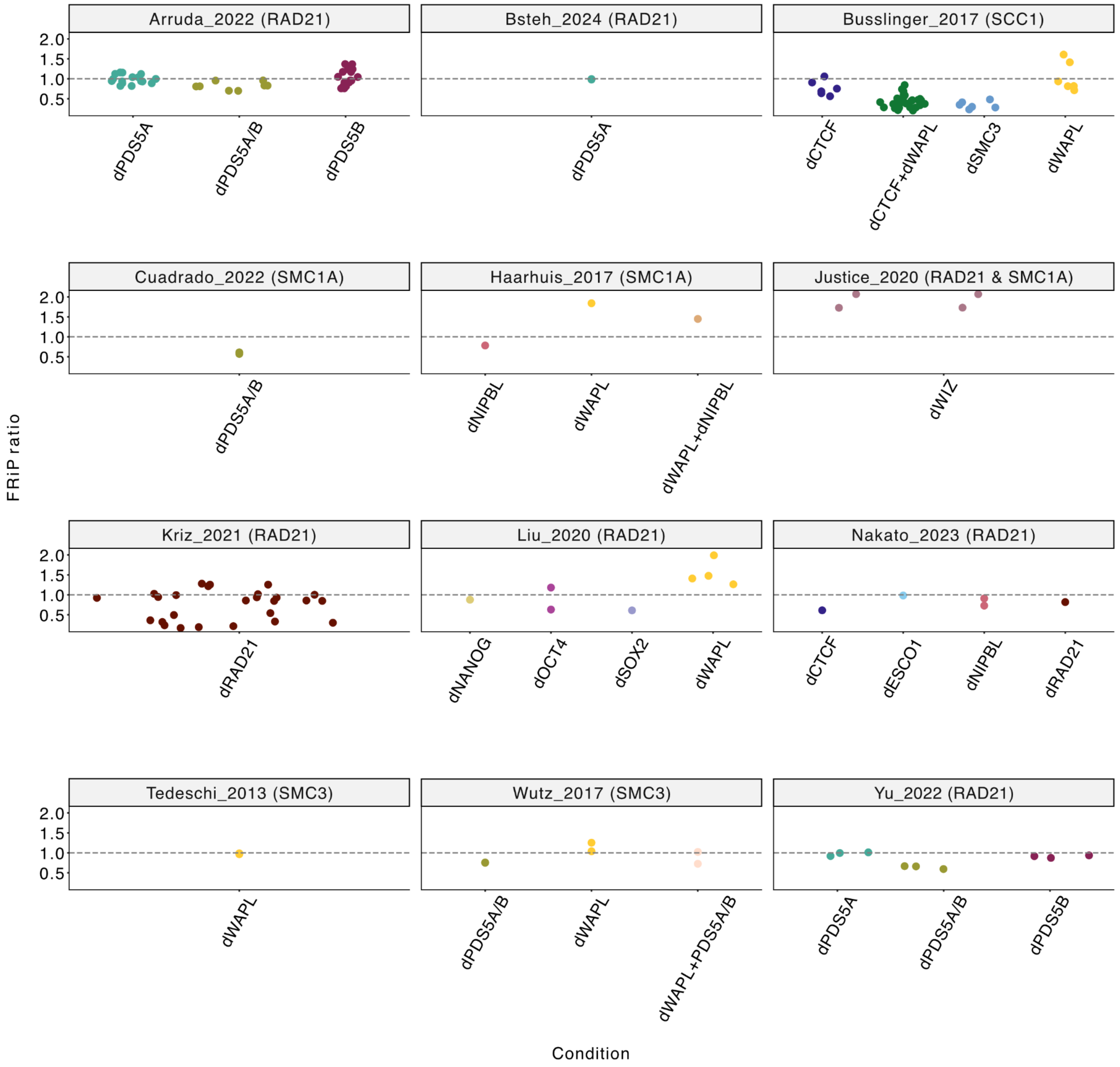


**Figure S5. FRiP ratios across studies.** 12 studies have conditions other than unperturbed. Each point represents an individual sample, with colors indicating experimental conditions. Y-axis shows a FRiP ratio of 1.0 indicates no change from unperturbed conditions. The “d” prefix denotes depletion of the corresponding cofactor.

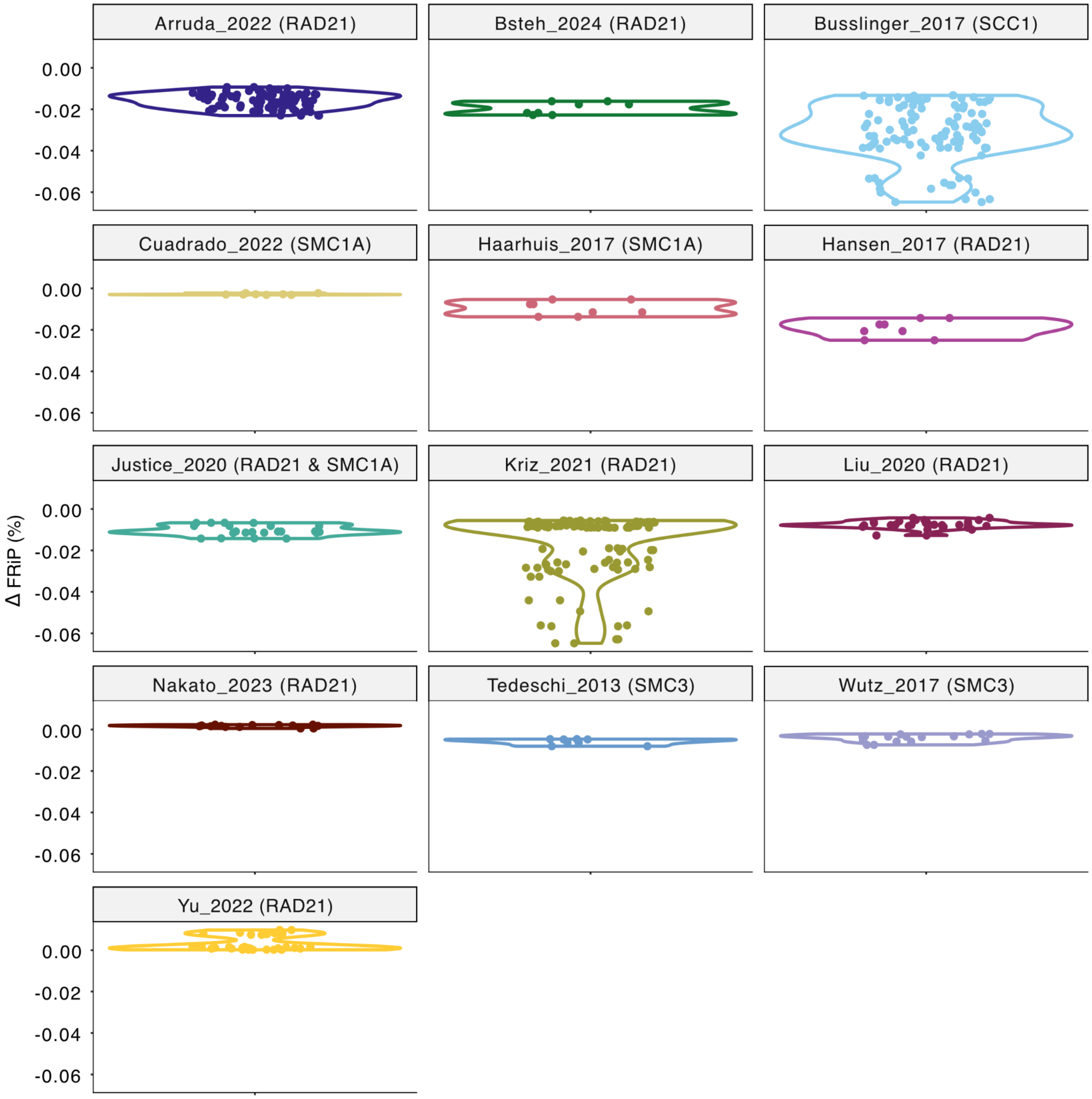

Arruda_2022 (RAD21)
Bsteh_2024 (RAD21)
Busslinger_2017 (SCC1)
Cuadrado_2022 (SMC1A)
Haarhuis_2017 (SMC1A)
Hansen_2017 (RAD21)
Justice_2020 (RAD21 & SMC1A)
Kriz_2021 (RAD21)
Liu_2020 (RAD21)
Nakato_2023 (RAD21)
Tedeschi_2013 (SMC3)
Wutz_2017 (SMC3)
Yu_2022 (RAD21)
Δ FRiP (%)
0.00
-0.02
-0.04
-0.06

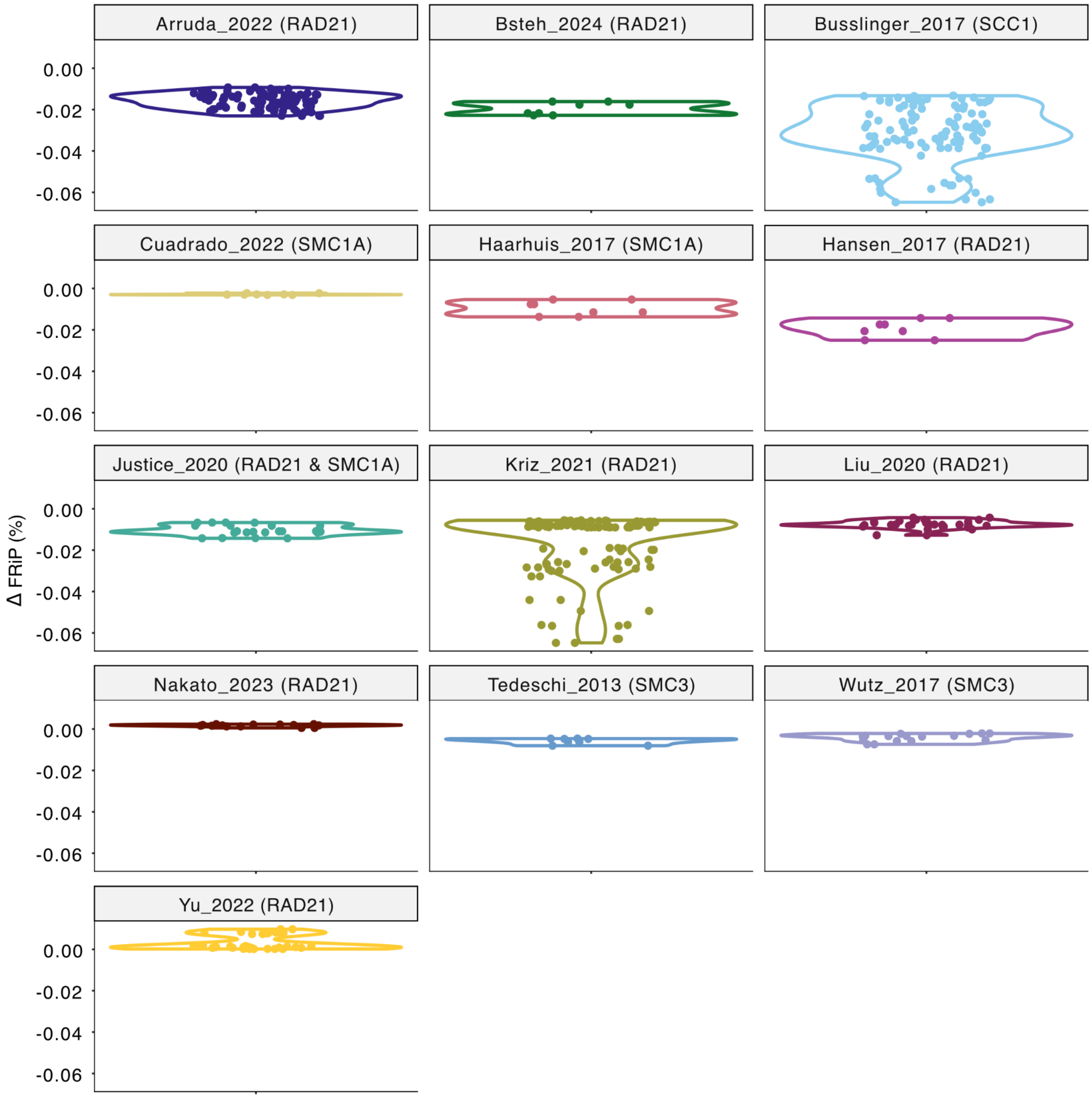


**Figure S6. Excluding blacklist regions has negligible impact on FRiP.** Each datapoint represents the proportional change in FRiP after excluding blacklist regions for each sample from the corresponding study, including both unperturbed and perturbed samples.

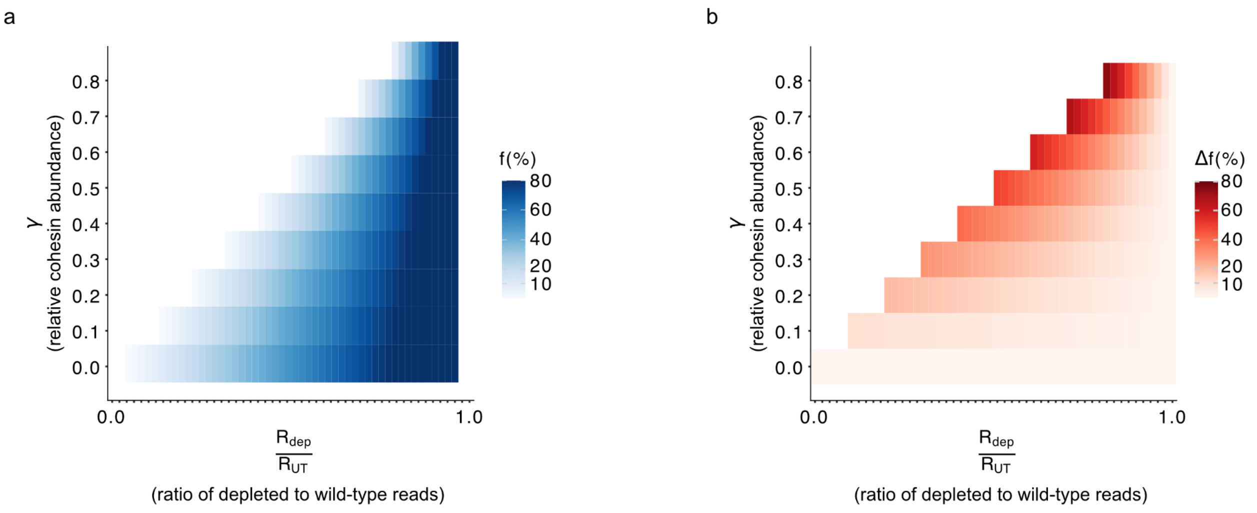


**Figure S7. Low depletion efficiency with high antibody quality results in large errors in background estimation. a)** heatmap of background fraction for various ratios of spike-in normalized reads and relative cohesin abundances, illustrating that the same read ratio can correspond to very different background fractions depending on how complete depletion is. **b)** heatmap of the error in estimated background fraction when complete depletion ($\gamma$ = 0) is assumed, showing that residual cohesin can substantially bias background estimation.

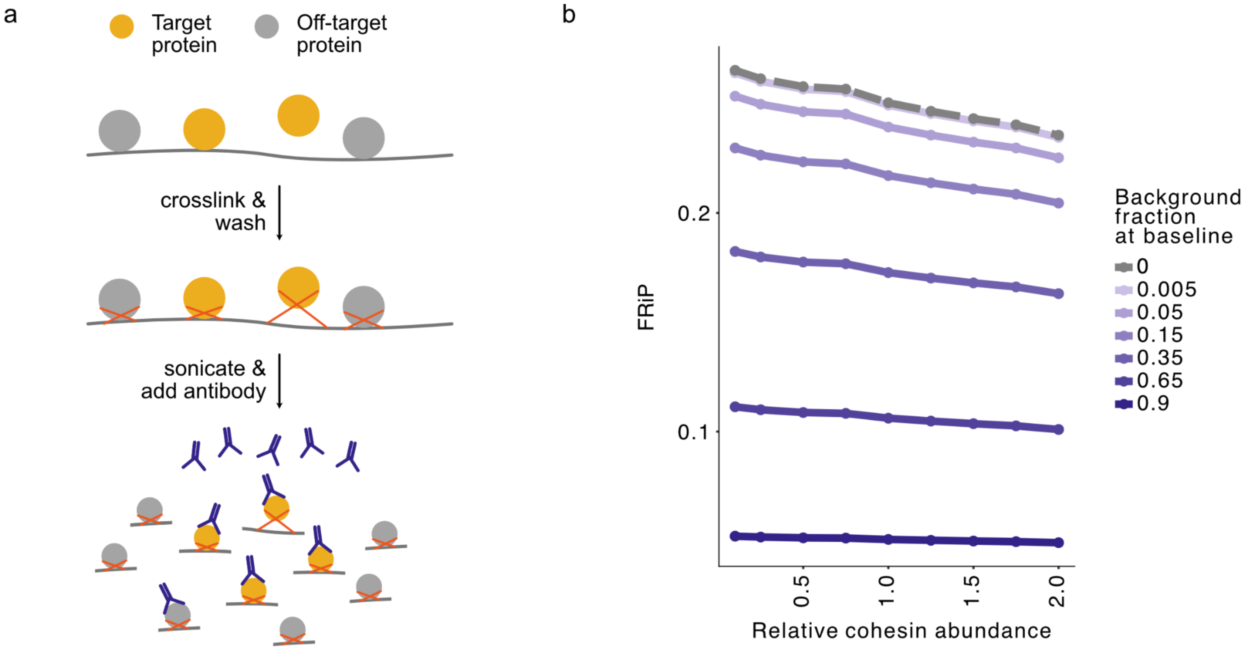


**Figure S8. Schematic of the alternative proximally-crosslinked ChIP model. a)** Formaldehyde randomly crosslinks target proteins proximal to chromatin even when they are not directly bound. With this model, antibodies capture both chromatin-bound and proximally-crosslinked proteins, both of which scale with target protein abundance. **b)** Under this model, FRiP decreases monotonously as cohesin abundance increases for any background fraction at baseline.

| | Author, Year | #Params | Support Input | Support no Input | Removes spike-in | Support no spike-in | Alignment software | Filtering software | Peak ca[illegible] software |
|---|---|---|---|---|---|---|---|---|---|
| **ChIP-FRiP** | **Xiao, 2025** | **12** | ✓ | ✓ | ✓ | ✓ | **Bowtie2** | **SAMtools** | **MACS2** |
| SpikeFlow | Bressan, 2024 | 39 | ✓ | X | ✓ | X | Bowtie2 | SAMtools | MACS2 |
| chipseq | Patel, 2024 | 53 | ✓ | X | X | ✓ | BWA / Chromap / Bowtie2 / STAR | SAMtools & BEDTools & BAMtool & Picard & Pysam | MACS3 |
| ENCODE | Landt, 2012 | 106 | ✓ | ✓ | X | ✓ | Bowtie2 | SAMtools & Picard | MACS2 |

**Table S1. Comparison of ChIP-seq pipelines.** While numerous high-quality ChIP-seq analysis pipelines exist, they have limitations for systematic cross-study analysis of cohesin regulation. For example, the ENCODE pipeline lacks support for spike-in normalization and uses a less widely-distributed workflow manager (Cromwell). SpikeFlow strictly requires both spike-in controls and input samples. This requirement precludes analysis of many publicly available datasets that provide only target protein ChIP-seq data. Furthermore, many pipelines offer extensive options for customization that, while providing flexibility, create barriers to usage as they require setting multiple parameters without clear guidance on optimal settings. For instance, SpikeFlow, chipseq, and ENCODE pipelines contain 39, 53, and 106 configuration parameters, respectively (#Params in the second column were derived from workflow configuration files or documentation). X indicates missing functionality for the specified feature. All pipelines mentioned here support both single-end and paired-end data.

| Author_year | GEO Accession | Species | cell type | is Spike-in | Antibody |
|---|---|---|---|---|---|
| Arruda_2022 | GSE199356 | Mus musculus | mESCs v6.5 | Yes | RAD21 (Abcam ab992) |
| Bsteh_2024 | GSE194268 | Mus musculus | mESCs | Yes | RAD21 (Abcam ab992); CTCF (Millipore 070729) |
| Busslinger_2017 | GSE76303 | Mus musculus | Mouse embryonic fibroblasts | No | scc1 (abcam ab992; Stag1 (Peters laboratory ID A823); CTCF (Upstate 07-729) |
| Cuadrado_2022 | GSE212151 | Mus musculus | Mouse embryonic fibroblasts | No | SMC1A (Custom made); CTCF (Millipore 07-729) |
| Haarhuis_2017 | GSE95015 | Homo sapiens | Hap1 | No | SMC1A (Bethyl, A300-055a); CTCF (Millipore, 07-72); |
| Hansen_2017 | GSE90994 | Mus musculus | JM8.N4 (mESCs) | No | Rad21 (Abcam ab154769); CTCF (Abcam ab128873) |
| Justice_2020 | GSE137285 | Mus musculus | mESCs v6.5 | Yes | anti-CTCF Active Motif 61311; anti-RAD21 Bethyl A300-080A; anti-SMC1A; Bethyl A300-055A |
| Kriz_2021 | GSE144834 | Mus musculus | mESCs (Tsix-stop) | Yes | RAD21 (Abcam ab992); CTCF (Active Motif 61311) |
| Liu_2020 | GSE135180 | Mus musculus | mESCs & Neural precursor | Yes | CTCF (07-729, Merck Millipore); RAD21 (ab154769, Abcam) |
| Nakato_2023 | GSE196450 | Homo sapiens | RPE | Yes | CTCF (07-729, Merck); Rabbit polyclonal antibody against Rad21; The mouse monoclonal antibody against the acetylated form of SMC3; |
| Tedeschi_2013 | GSE41603 | Mus musculus | Mouse embryonic fibroblasts | No | Smc3 (Bethyl A300-060A); CTCF (Upstate 07-729) |
| Wutz_2017 | GSE102884 | Homo sapiens | HeLa (Kyoto) | No | Anti-CTCF Merck Milipore Cat# 07–729; Anti-SMC3 Peters laboratory Antibody ID:A941; Anti-STAG1 Peters laboratory Antibody ID:A823 |
| Yu_2022 | GSE209849 | Homo sapiens | PLC/PRF-5 (hepatoma) | Yes | ctcf (millipore, 07-729); rad21 (abcam, ab992) |

**Table S2. Study metadata.** Studies are identified by the first author name and publication year.

**Supplementary note**

***Derivation of the biochemical model of background noise***

To arrive at Equation 2, we modeled antibody (A) binding to the chromatin-bound cohesin (C) and background (B) with two equilibrium binding equations:

$$[A][C] = K_d{}^s[AC] \quad \text{(Eq. S1)}$$

$$[A][B] = K_d{}^n[AB] \quad \text{(Eq. S2)}$$

Where $K_d{}^s$ is the dissociation constant for specific binding, and $K_d{}^n$ is the dissociation constant for non-specific binding. We also have three equations for the total number of antibodies ($A^{total}$), cohesin ($C^{total}$), and background proteins ($B^{total}$):

$$[A^{total}] = [A] + [AC] + [AB] \quad \text{(Eq. S3)}$$

$$[B^{total}] = [B] + [AB] \quad \text{(Eq. S4)}$$

$$[C^{total}] = [C] + [AC] \quad \text{(Eq. S5)}$$

Here, $[A]$ is the concentration of free antibody, $[B]$ is the concentration of free background proteins, $[C]$ is the concentration of loaded cohesins that are not bound by antibodies. Substituting the expression for $[C^{total}]$ into the equilibrium equation for $[AC]$:

$$[AC] = \frac{[C^{total}]}{\frac{K_d{}^s}{[A]} + 1} = [C^{total}] * \alpha \quad \text{(Eq. S6)}$$

Where $\alpha$ represents the proportion of $[C^{total}]$ bound by antibodies.

Following Abcam's ChIP assay guidelines(50), we assumed reference values of: chromatin corresponding to 25μg DNA, 5μg antibodies, and 500μL reaction volume.

To estimate the number of 1Mb sequences in this reaction volume, we used an average molecular weight of a single DNA base pair as 650 g/mol. Thus, 25μg DNA contains around $2.3161538461 * 10^{10}$ sequences of 1Mb. We then transformed the unit of cohesin abundance to molarity with the scale transformation constant:

$$\sigma = \frac{Counts\ of\ 1Mb\ sequences\ in\ reaction\ /\ N_A}{Volume\ of\ IP\ reaction} = \frac{23161538461\ /(6.022 * 10^{23})}{5 * 10^{-4}\ L} \quad \text{(Eq. S7)}$$

By multiplying the expected number of loaded cohesins per 1Mb (4) by $\sigma$, we obtain an estimated concentration of loaded cohesin $[C^{total}]$ before ChIP-seq, $3.077 * 10^{-10}\ M$. We used the molecular weight of an IgG antibody (150000 g/mol) (51) to calculate $[A^{total}]$: $6.66 * 10^{-8}\ M$ in 500μL.

From these literature parameters, $[A^{total}]$ is much higher than $[C^{total}]$ (0.46% of $[A^{total}]$). For background fractions up to 90% (i.e. a nine-fold excess over cohesin reads) $[AB]$ is thus less than 4.15% of $[A^{total}]$. ChIP-seq experiments with higher background fractions likely lack interpretability. Since $[AC]$ is less than $[C^{total}]$, we can thus approximate Equation S3 as: $[A^{total}] \approx [A]$.

To understand how $[AB]$ relates to $[AC]$, we substituted the expression for $[B^{total}]$ into the equilibrium equation for [AB]:

$$[AB] = \frac{[B^{total}]}{\frac{K_d{}^n}{[A]} + 1} = [A] * \frac{[B^{total}]}{K_d{}^n + [A]} = [A] * \beta \approx [A^{total}] * \beta \qquad \text{(Eq. S8)}$$

Where we defined $\frac{[B^{total}]}{K_d{}^n + [A]}$ as the non-specific binding ratio $\beta$.
With the approximation for $[A^{total}]$, $[AB]$ is independent of $[AC]$ (or $[C]$ or $[C^{total}]$). Given $[AB]$ is independent of the amount of cohesin, we can simply define $[AB] * reaction\ volume + \epsilon$ as a constant $\theta$, representing total background reads, and arrive at Equation 2:

$$Total\ ChIP\ reads \propto [AC] + \theta$$

***Derivation of the modified background fraction***
We defined $f$ as the following expression:

$$f = \frac{\theta}{[AC] + \theta} \qquad \text{(Eq. S9)}$$

We formulated $\theta$ as a function of $f_{base}$:

$$\theta = \frac{f_{base}}{1 - f_{base}} * [AC] \qquad \text{(Eq. S10)}$$

Substituting Equation S10 into Equation S9, we arrive at an expression for the modified background fraction:

$$modified\ background\ fraction = f_{mut}(\gamma) = \frac{\frac{f_{base}}{1 - f_{base}} * [AC]}{[AC] * \gamma + \frac{f_{base}}{1 - f_{base}} * [AC]} = \frac{f_{base}}{\gamma\,(1 - f_{base}) + f_{base}}$$

***Derivation of the background fraction of the unperturbed sample***
After depletion, background reads are $f_{dep} * R_{dep}$ with $f_{dep}$ the background fraction in the depletion sample. Similarly, remaining cohesin reads can be written in terms of the original background fraction and the degree of depletion:

$$Remaining\ Cohesin\ Reads = \gamma * UT\ Cohesin\ Reads = \gamma * (1 - f_{UT}) * R_{UT} \qquad \text{(Eq. S11)}$$

Where $\gamma$ represents the relative cohesin abundance, $f_{UT}$ is the background fraction of the unperturbed sample.

Using Equations 3 and S11, we substitute into Equation 4 to derive:

$$R_{dep} \quad = \frac{f_{UT}}{\gamma\,(1 - f_{UT}) + f_{UT}} * R_{dep} \quad + \gamma * (1 - f_{UT}) * R_{UT} \qquad \text{(Eq. S12)}$$

Rearranging Equation S12 to solve for $f_{UT}$ and arrive at Equation 5.

***Proximally-crosslinked ChIP model***

An alternative model that includes proximally-crosslinked proteins is presented in Fig. S8. In this model, a fraction $\delta$ of $[AC]$ derives from proximally crosslinked cohesins rather than chromatin-bound cohesins. Thus, proximally crosslinked cohesins constitute a second source of background reads, distinct from non-specific antibody binding.

$$Total\ ChIP\ reads \ \propto \ (1 - \delta) * [AC] \ + \theta \ + \ \delta * [AC]$$

We modified the following equations to account for this updated background composition:

$$f \quad = \frac{\theta + \delta * [AC]}{[AC] + \theta} \qquad \text{(Eq. S13)}$$

$$\theta = \frac{(f_{base} - \delta)}{1 - f_{base}} \ * \ [AC] \qquad \text{(Eq. S14)}$$

$$f_{mut}(\gamma) \ = \frac{\frac{f_{base} * (1 - \delta)}{1 - f_{base}} * [AC]}{[AC] * \gamma + \frac{f_{base} * (1 - \delta)}{1 - f_{base}} * [AC]} \ = \frac{f_{base} * (1 - \delta)}{\gamma\,(1 - f_{base}) + f_{base} * (1 - \delta)} \qquad \text{(Eq. S15)}$$

$$R_{dep} \quad = \frac{f_{UT} \ * (1 - \delta)}{\gamma\,(1 - f_{UT}) + f_{UT} * (1 - \delta)} * R_{dep} \quad + \gamma * (1 - f_{UT}) * R_{UT} \qquad \text{(Eq. S16)}$$

$$f_{UT} \ = \frac{\frac{R_{dep}}{R_{UT}} - \gamma}{1 - \gamma - \delta} \qquad \text{(Eq. S17)}$$

If we set $\delta \ = \ 0$, then the equations are the same as our primary model.
